# Development of *Arabidopsis thaliana* transformants showing the self-recognition activity of *Brassica rapa*

**DOI:** 10.1101/2020.07.15.205708

**Authors:** Masaya Yamamoto, Hiroyasu Kitashiba, Takeshi Nishio

## Abstract

Self-incompatibility in the Brassicaceae family is governed by two-linked highly polymorphic genes located at the *S* locus, *SRK* and *SCR*. Previously, the *SRK* and *SCR* genes of *Arabidopsis lyrata* were introduced into *Arabidopsis thaliana* transformants to generate self-incompatible lines. However, it has not been reported that the *SRK* and *SCR* genes of *Brassica* species confer self-incompatibility in *A. thaliana*. In this study, we attempted to construct self-incompatible *A. thaliana* transformants expressing the self-recognition activity of *Brassica rapa* by introducing the *BrSCR* gene along with a chimeric *BrSRK* gene (*BrSRK chimera*, in which the kinase domain of *BrSRK* was replaced with that of *AlSKRb*). We found that *BrSRK chimera* and *BrSCR* of *B. rapa S-9* and *S-46* haplotypes, but not those of *S-29*, *S-44*, and *S-60* haplotypes, conferred self-recognition activity in *A. thaliana*. We also investigated the importance of amino acid residues involved in the BrSRK9–BrSCR9 interaction using *A. thaliana* transformants expressing mutant variants of *BrSRK-9 chimera* and *BrSCR-9*. The results showed that some of the amino acid residues are essential for self-recognition. The method developed in this study for the construction of self-incompatible *A. thaliana* transformants showing *B. rapa* self-recognition activity will be useful for analysis of self-recognition mechanisms in Brassicaceae.

## INTRODUCTION

Self-incompatibility (SI) is one of the key strategies used to promote outcrossing and maintain genetic diversity in flowering plants. In the Brassicaceae family, SI is governed by two polymorphic genes located at the *S* locus: *S-locus receptor kinase* (*SRK*) and *S-locus cysteine-rich protein*/*S-locus protein 11* (*SCR*/*SP11*; hereafter referred to as *SCR*). Because alleles of *SRK* and *SCR* are inherited as a set, this set is called the *S* haplotype. The *SRK* gene encodes plasma membrane-localized serine/threonine (Ser/Thr) receptor kinase expressed in stigma papillar cells [1], while the *SCR* gene encodes pollen-surface localized SRK ligand [2–4]. SRK interacts with SCR in an allele-specific manner [3, 4]. The binding of SRK with SCR of the same *S* haplotype activates SI, which inhibits the germination of self-pollen and elongation of the self-pollen tube in stigma.

Historically, most of the SI research in Brassicaceae has been carried out using the *Brassica* species. Biochemical analysis of SI in *Brassica* has contributed to the elucidation of allele-specific interaction between SRK and SCR [3, 4]. SCR typically has eight conserved cysteine (Cys) residues, and experiments using recombinant *Brassica* SCR proteins show that two amino acid regions (one between the second and third Cys residues, and the other between the fifth and sixth Cys residues) determine the recognition specificity of SCR [5, 6]. Moreover, studies using interspecific hybrids between *Brassica rapa* and *Brassica oleracea* indicate that several *S* haplotypes have the same recognition specificities between these two species [6, 7]. Recently, the structure of the BrSRK9 ectodomain (hereafter referred to as eBrSRK9) and BrSCR9 complex was solved, which revealed the amino acid residues involved in the eBrSRK9–BrSCR9 interaction [8].

The model dicot plant *Arabidopsis thaliana*, which belongs to the family Brassicaceae, does not exhibit SI because it contains a nonfunctional *SCR* gene or both nonfunctional *SCR* and *SRK* genes [9–13]. The *SRK* and *SCR* genes of self-incompatible species closely related to *A. thaliana*, including *Arabidopsis lyrata*, *Arabidopsis halleri*, and *Capsella grandiflora*, confer SI in *A. thaliana* [14–18]. These findings indicate that *A. thaliana* possesses all components required to activate SI, except SCR (and SRK). However, since production of self-incompatible *A. thaliana* plants by introducing *Brassica SRK* and *SCR* genes has been unsuccessful [19], *in planta* evaluation of the results of biochemical and structural studies on these two *Brassica* genes has been limited.

Recent genomic analyses revealed the structure of the *S* locus in several self-incompatible and -compatible Brassicaceae species [20–22]. In all Brassicaceae genera, except *Brassica* and *Leavenworthia*, *SRK* and *SCR* (or their pseudogenes) are flanked by *PUB8* (*At4g21350*) and *ARK3* (*At4g21380*), and are located in block U of the ancestral Brassicaceae karyotype [23]. In *B. rapa*, *BrSRK* and *BrSCR* genes are flanked by *BrSP2* and *BrSP6* [24] (*At1g66670* and *At1g66730*, respectively), and are located in block E of the ancestral Brassicaceae karyotype [25]. By contrast, no gene is located between *Bra013521* and *Bra013522*, which are orthologous to *AtPUB8* and *AtARK3*, respectively. In addition to the difference at the *S* locus, *A. thaliana* does not harbor genes orthologous to the *Brassica* SI modifier genes, such as *MLPK* and *ARC1* [26–29]. Loss-of-function mutations in *AtAPK1b* and *AtPUB17*, which are most similar to *Brassica MLPK* and *ARC1*, respectively, do not affect the SI trait of the self-incompatible *A. thaliana* transformants [26, 30]. These reports suggest the possibility that *Brassica* and *Arabidopsis* possess distinct SI signaling cascades.

More than 40 *S* haplotypes have been identified in *B. rapa* [31, 32], and they are divided into two groups, class I and class II, based on sequence similarity. In this study, to generate *A. thaliana* plants showing the *Brassica* self-recognition specificity, we constructed *thaliana* transformants expressing the *BrSRK chimera*, which possesses the *S* domain and transmembrane domain of *BrSRK*, followed by the kinase domain of *AlSRKb*, and *BrSCR* genes of two class-I (*S-9*, and *S-46*) and three class-II (*S-29*, *S-44*, and *S-60*) haplotypes. We found that the *A. thaliana* transformants expressing the *BrSRK-9 chimera* and *BrSCR-9* (*BrSRK-9 chimera*+*BrSCR-9*) and *BrSRK-46 chimera*+*BrSCR-46* showed SI.

Self-incompatible *A. thaliana* transformants expressing *BrSRK-9 chimera*+*BrSCR-9* were further used to identify the essential amino acid residues of BrSRK9 and BrSCR9, which have been reported to be involved in the eBrSRK9–BrSCR9 interaction [8], for SI. Thus, the method developed in this study, which allows the generation of self-incompatible *A. thaliana* transformants, can be used to analyze the mechanisms of recognition specificities in Brassicaceae.

## RESULTS

### BrSRK9 kinase domain does not function in *A. thaliana*

To explore why *Brassica SRK* and *SCR* fail to induce SI in *A. thaliana*, we constructed *A. thaliana* transformants expressing *BrSRK-9* under the control of the *AtS1* promoter, which is a highly active, stigma papillar cell-specific promoter of *A. thaliana* [33], and *BrSCR-9* under the control of the *AlSCR-b* promoter; these *A. thaliana* transformants are hereafter referred to as *BrSRK-9*+*BrSCR-9* (Figure 1A). Flower buds just before flowering (−1 position) of *A. thaliana* transformants were self-pollinated, and the number of pollen tubes per pollinated stigma was counted. More than 30 elongating pollen tubes were observed in the stigmas of all tested *BrSRK-9*+*BrSCR-9 A. thaliana* transformants (Figure 1B, Table 1), which was similar to the observation in untransformed wild-type *A. thaliana* plants (Figure 1B).

**Table 1.** Self-incompatibility (SI) phenotypes of *Arabidopsis thaliana* transformants expressing *Brassica rapa S-9* genes.

| Transgenes | Total no. of transformants | No. of self-pollen tubes per stigma<sup>a</sup> |  |  |
| --- | --- | --- | --- | --- |
|  |  | <10 | 10–29 | >29 |
| *BrSRK-9+BrSCR-9* | 13 | 0 | 0 | 13 |
| *BrSRK-9 chimera+BrSCR-9* | 14 | 10 | 2 | 2 |
| *BrSRK-9+BrSCR-9+MLPK* | 13 | 0 | 0 | 13 |

<sup>a</sup>The number of pollen tubes per stigma was counted at 2 h after self-pollination. Values of <10, 10–29, and >29 indicate strong SI, weak SI, and compatible responses, respectively.

**Figure 1.**
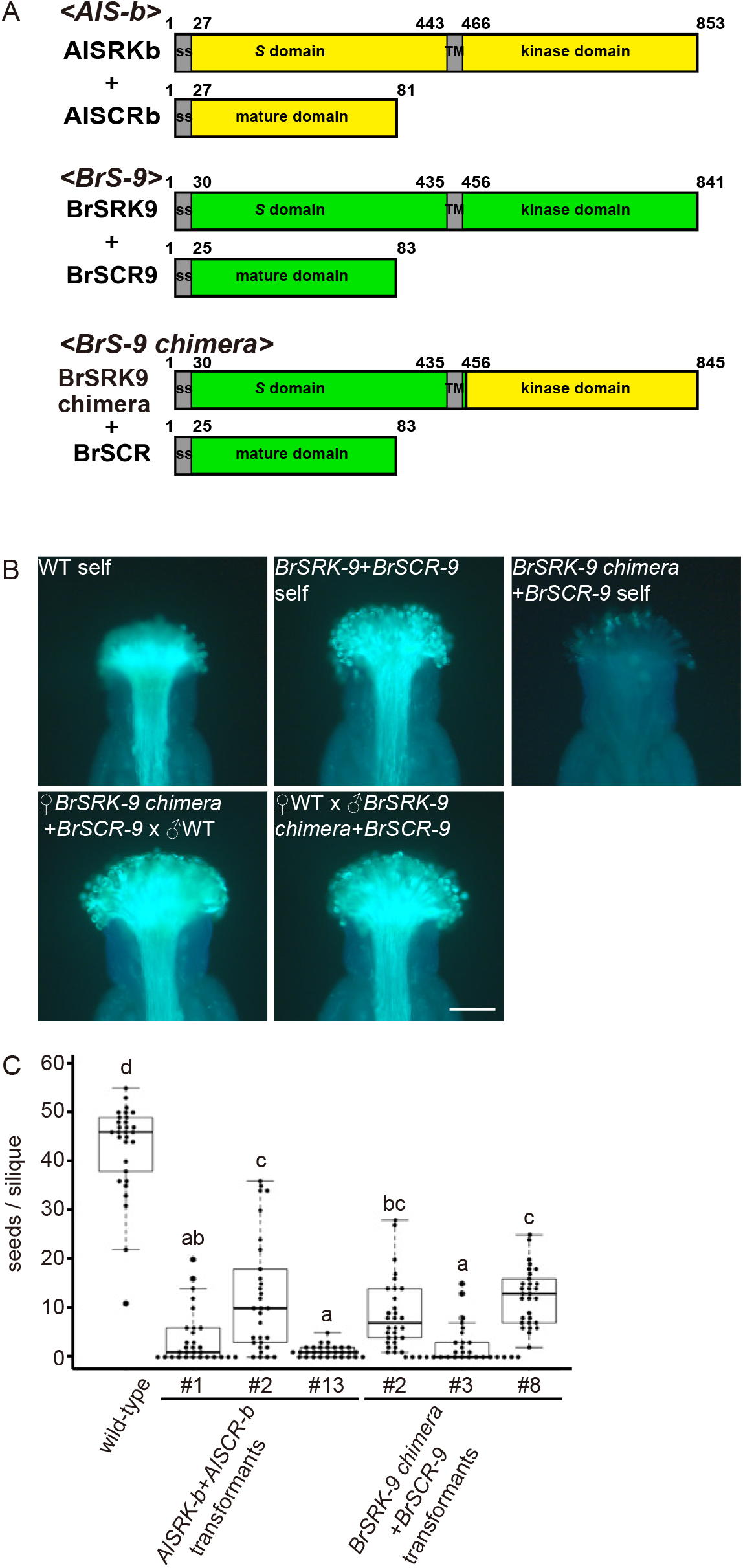
Effects of *BrSRK-9*+*BrSCR-9* and *BrSRK-9 chimera*+*BrSCR-9* on self-incompatibility (SI) in *Arabidopsis thaliana* plants. (A) Schematic showing the structures of AlSRKb and AlSCRb (top), BrSRK9 and BrSCR9 (middle), and BrSRK9 chimera and BrSCR9 (bottom). The kinase domain of BrSRK9, just after the transmembrane region, was replaced with that of AlSRKb, and the site where the kinase domain of BrSRK was replaced by that of AlSRKb in BrSRK chimeras is shown in Supplemental Figure 3. Al, *Arabidopsis lyrata*; Br, *Brassica rapa*. (B) Images showing self-pollinated (upper panel) and cross-pollinated (lower panel) stigmas of *A. thaliana* plants stained with aniline blue. WT self: self-pollinated stigma of wild-type plants; *BrSRK-9*+*BrSCR-9* self: self-pollinated stigma of *BrSRK-9*+*BrSCR-9* transgenic plants; *BrSRK-9 chimera*+*BrSCR-9* self: self-pollinated stigma of *BrSRK-9 chimera*+*BrSCR-9* transgenic plants; ♀*BrSRK-9 chimera*+*BrSCR-9* × ♂WT: stigma of the *BrSRK-9 chimera*+*BrSCR-9* transgenic plant cross-pollinated with wild-type pollen; ♀WT x ♂*BrSRK-9 chimera*+*BrSCR-9*: wild-type stigma cross-pollinated with *BrSRK-9 chimera*+*BrSCR-9* transgenic pollen. Scale bar = 100 μm. (C) Box-plot analysis of the number of seeds per silique of wild-type plants, *AlSRK-b*+*AlSCR-b* transformants, and *BrSRK-9 chimera*+*BrSCR-9* transformants of *A. thaliana*. Data were obtained from 30 siliques per plant. Different letters indicate significant differences (*P* < 0.05; Tukey–Kramer method).

It could be argued that the *BrSRK-9* and *BrSCR-9* genes are nonfunctional in *A. thaliana* because of the atypical expression of *BrSRK-9* and *BrSCR-9*. However, to exclude the possibility, we examined the expression of *BrSRK-9* and *BrSCR-9* in the buds of the constructed *A. thaliana* transformants. First, to determine the flower/bud developmental stages for expression analysis of the *SRK* and *SCR* genes, we analyzed *SRK* and *SCR* expression during flower development in the self-incompatible *AlSRK-b*+*AlSCR-b A. thaliana* transformants, which express *AlSRK-b* and *AlSCR-b* under the control of *AtS1* and *AlSCR-b* promoters, respectively [34]. We divided flower development into four stages (from old to young): immediately after and at flowering (+1 and 0 positions, respectively), immediately before flowering and the second bud before flowering (−1 and −2, respectively), the third and fourth buds before flowering (−3 and −4, respectively), and the fifth and sixth buds before flowering (−5 and −6, respectively). Given that *AlSRK-b* transcript levels showed no differences among the tested flowers and buds at different developmental stages (Supplemental Figure 1, Supplemental Table 1), and the fact that buds at position −1 were used for the pollination assay, we analyzed *SRK* expression levels in buds at positions −1 and −2. Since buds at positions −3 and −4 showed the highest level of *AlSCR-b* transcripts (Supplemental Figure 1, Supplemental Table 1), we used these buds for *SCR* expression analysis in subsequent experiments.

Absolute quantitative real-time PCR (qRT-PCR) analysis showed that the expression of *BrSRK-9* in the buds (positions −1 and −2) of three *BrSRK-9*+*BrSCR-9* transgenic *A. thaliana* lines were higher than that of *AlSRK-b* in the corresponding buds of three *AlSRK-b*+*AlSCR-b* transgenic and self-incompatible *A. thaliana* lines [34] (Supplemental Figure 2, Supplemental Table 2), indicating that the nonfunctionality of *BrSRK-9* in *A. thaliana* is not due to reduced transcription levels of *BrSRK-9*. Although transcript levels of *BrSCR-9* were lower than those of *AlSCR-b* in the three *AlSRK-b*+*AlSCR-b* transgenic *A. thaliana* lines (Supplemental Figure 2, Supplemental Table 3), the *BrSCR-9* gene was expressed in all tested *BrSRK-9+BrSCR-9* transgenic *A. thaliana* lines.

Next, we constructed *BrSRK9 chimera*+*BrSCR9* transgenic *A. thaliana* plants (Figure 1A, Supplemental Figure 3). In the self-pollination assay, no or few pollen tubes were observed in the stigmas of the 10 constructed *BrSRK-9 chimera*+*BrSCR-9* transgenic *A. thaliana* lines (Figure 1B, Table 1). A large number of wild-type pollen tubes were observed in the stigmas of *BrSRK-9 chimera*+*BrSCR-9* transgenic *A. thaliana* plants and vice versa (Figure 1B), indicating that the pollen and stigmas of the constructed transformants functioned normally. We then compared the number of seeds per silique produced by the autonomously self-pollinating flowers of different genotypes. The results showed that the number of seeds per silique in three *BrSRK-9 chimera*+*BrSCR-9* transgenic *A. thaliana* lines was lower than that in wild-type plants (Figure 1C), but did not differ significantly from that in three *AlSRK-b*+*AlSCR-b* transgenic *A. thaliana* lines. These results indicate that *BrSRK-9 chimera*+*BrSCR-9* confer SI in *A. thaliana*.

Since knockout mutation of the *MLPK* gene in self-incompatible *Brassica napus* lines using the CRISPR/Cas9 technology caused defects in SI [35], and MLPK proteins in *Escherichia coli* were phosphorylated by recombinant SRK kinase domain proteins [36], MLPK is considered as a substrate of SRK and to function in SI. Therefore, to examine whether the expression of *BrMLPK*, *BrSRK-9*, and *BrSCR-9* in *A. thaliana* plants causes SI, we introduced the *AtS1*_*pro*_:*BrMLPK* construct into *BrSRK-9*+*BrSCR-9 A. thaliana* line #1, resulting in 13 transformants. A large number of self-pollen tubes were observed in the stigmas of all *BrSRK-9*+*BrSCR-9*+*BrMLPK* transgenic *A. thaliana* lines (Supplemental Figure 4A, Table 1). In addition to *BrSRK-9* transcripts, *BrMLPK* transcripts were detected in the buds (positions −1 and −2) of *BrSRK-9*+*BrSCR-9*+*BrMLPK A. thaliana* lines #1, #2, and #3 by reverse transcription PCR (RT-PCR) (Supplemental Figure 4B). These results indicate that co-expression of *BrSRK-9*, *BrSCR-9*, and *BrMLPK* is not sufficient for inducing SI in *A. thaliana*.

### *B.rapa S-9* and *S-46* haplotypes, but not *S-29*, *S-44*, and *S-60* haplotypes, confer self-recognition activity in *A. thaliana*

To examine whether other *S* haplotypes of *B. rapa* confer self-recognition activity in *A. thaliana*, we tested the SI of *A. thaliana* transformants expressing the *BrSRK chimera*+*BrSCR* genes of *S-46*, *S29*, *S-44*, and *S-60* haplotypes. In the manual pollination assay, no or few self-pollen tubes were observed in the stigmas of 9 out of 16 *BrSRK-46 chimera*+*BrSCR-46* transgenic *A. thaliana* lines (Figure 2A, Table 2). We also examined the number of seeds per silique of three *BrSRK-46 chimera*+*BrSCR-46* transgenic *A. thaliana* lines (#2, #7, and #10), which showed SI in the pollination assay. The number of seeds per silique of these lines was statistically lower than that observed in wild-type plants (Figure 2B), indicating that *BrSRK-46 chimera*+*BrSCR-46* confer SI in *A. thaliana*.

**Table 2.** SI phenotypes of transgenic *A. thaliana* plants co-expressing *BrSRK* and *BrSCR* genes.

| Transgenes | Total no. of transformants | No. of self-pollen tubes per stigma<sup>a</sup> |  |  |
| --- | --- | --- | --- | --- |
|  |  | <10 | 10–29 | >29 |
| *BrSRK-46 chimera+BrSCR-46* | 16 | 9 | 2 | 5 |
| *BrSRK-29 chimera+BrSCR-29* | 13 | 0 | 0 | 13 |
| *BrSRK-44 chimera+BrSCR-44* | 12 | 0 | 0 | 12 |
| *BrSRK-60 chimera+BrSCR-60* | 15 | 0 | 0 | 15 |

<sup>a</sup>The number of pollen tubes per stigma was counted at 2 h after self-pollination. Values of <10, 10–29, and >29 indicate strong SI, weak SI, and compatible responses, respectively.

**Figure 2.**
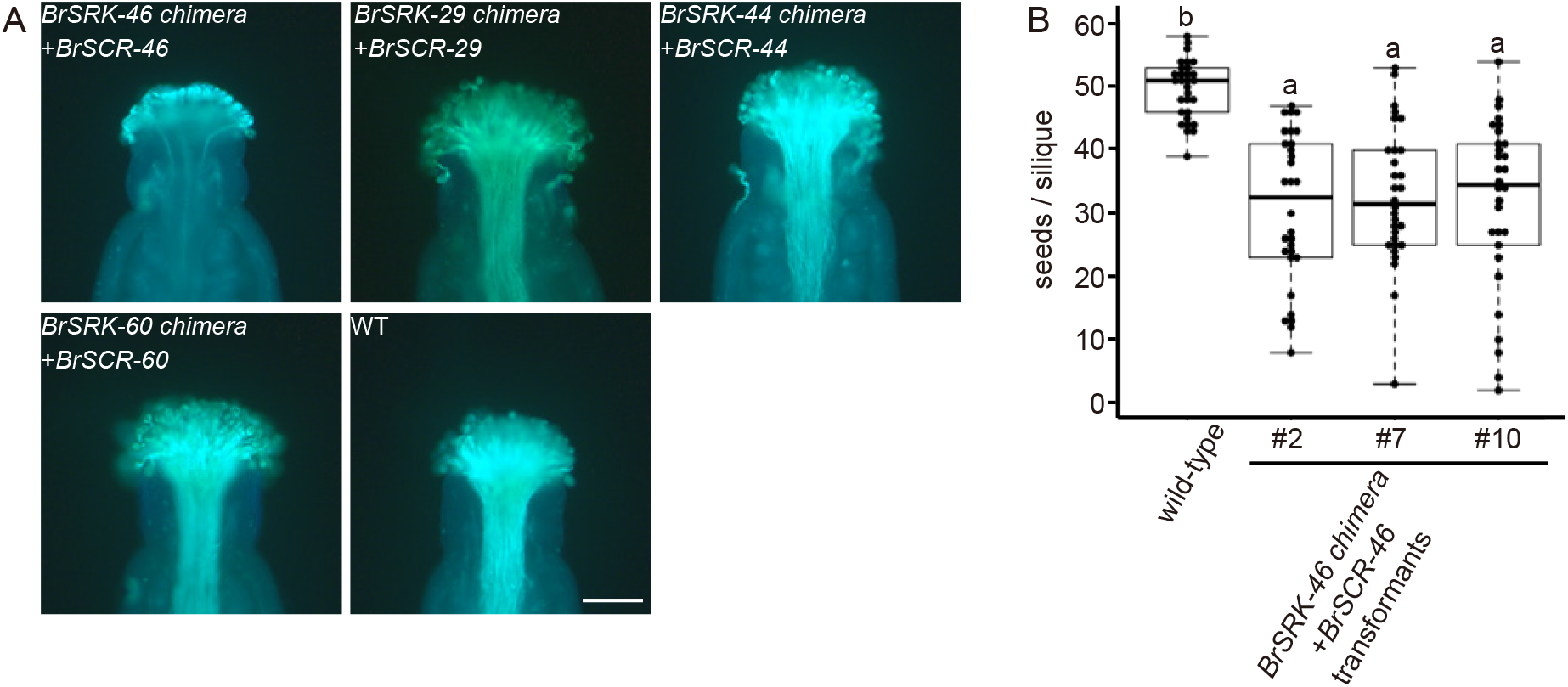
Functional analysis of *BrSRK chimera*+*BrSCR* of different *S* haplotypes in *A. thaliana*. (A) Representative images showing self-pollinated stigmas of wild-type (WT) and transgenic *A. thaliana* plants. Scale bar = 100 μm. (B) Box-plot analysis of the number of seeds per silique of wild-type plants and *BrSRK-46 chimera*+*BrSCR-46* transformants (*n* = 30 per plant). Different letters indicate significant differences (*P* < 0.05; Tukey–Kramer method).

On the other hand, *A. thaliana* transformants expressing *BrSRK-29 chimera*+*BrSCR-29*, *BrSRK-44 chimera*+*BrSCR-44*, and *BrSRK-60 chimera*+*BrSCR-60* allowed the penetration of numerous self-pollen tubes into the stigmas, similar to that observed in wild-type plants (Figure 2A, Table 2). We examined the expression levels of *BrSRK chimera* and *BrSCR* in the constructed transformants by absolute qRT-PCR analysis. In the buds (positions −1 and −2) of the constructed transformants, transcript levels of *BrSRK-29 chimera*, *BrSRK-44 chimera*, and *BrSRK-60 chimera* were not significantly different from those of *BrSRK-9 chimera* and *BrSRK-46 chimera* (Supplemental Figure 2, Supplemental Table 2). Similarly, *BrSCR* transcript levels showed no significant differences among the constructed *A. thaliana* lines expressing *BrSRK chimera*+*BrSCR* of different *S* haplotypes (Supplemental Figure 2, Supplemental Table 3). These results indicate that the pairs *BrSRK-29 chimera* and *BrSCR-29*, *BrSRK-44 chimera* and *BrSCR-44*, and *BrSRK-60 chimera* and *BrSCR-60* do not function in *A. thaliana*.

### Functional analysis of amino acid residues involved in the eBrSRK9–BrSCR9 *in planta*

Since amino acid residues involved in the eBrSRK9–BrSCR9 interaction have been predicted based on the structure of the eBrSRK9–BrSCR9 complex [8], we verified the role of some of these residues in activating SI. It has been reported that the short hairpin loop located between the second and third β-strands of BrSCR9 interacts with a hydrophobic cavity formed by five amino acid residues of BrSRK9 in the eBrSRK9–BrSCR9 complex, including valine at position 211 (V211), phenylalanine at positions 267 and 290 (F267 and F290, respectively), and proline at positions 287 and 294 (P287 and P294, respectively) [8]. Phenylalanine at position 69 (F69) of BrSCR9 has been reported to be the center of this interface, while tyrosine at position 73 (Y73) participates in interactions at the interface [8]. Therefore, we constructed *BrSRK-9 chimera*+*BrSCR-9(F69E)* and *BrSRK-9 chimera*+*BrSCR-9(Y73L)* transgenic *A. thaliana* plants (Figure 3A), and mutated four of the five residues (V211, P287, F290, and P294) of BrSRK9 that form the hydrophobic cavity in the eBrSRK9–BrSCR9 complex (Figure 3A). SI activity was tested by monitoring self-pollen germination and tube elongation. More than 30 self-pollen tubes were observed in the stigmas of these transgenic *A. thaliana* lines (Figure 3B, Table 3), indicating that these amino acid substitutions cause a defect in SI. The results of *in planta* analysis of *A. thaliana* transformants expressing *BrSRK-9(V211E) chimera*+*BrSCR-9*, *BrSRK-9(P294M) chimera*+*BrSCR-9*, and *BrSRK-9 chimera*+*BrSCR-9(F69E)* were consistent with those of *in vitro* analysis, where BrSRK9(V211E), BrSRK9(P294M), and BrSCR9(F69E) mutant proteins showed a defect in eBrSRK9–BrSCR9 interaction in the gel filtration assay [8]. We also examined the role of lysine at position 206 (K206) of BrSRK9 in activating SI. K206 of BrSRK9 is reportedly located at the edge of the eBrSRK9–BrSCR9 interface and interacts with proline at position 68 (P68) of BrSCR9 [8]. The results showed that 10 out of 17 *BrSRK-9(K206L) chimera*+*BrSCR-9* transgenic *A. thaliana* lines inhibited self-pollen germination and tube elongation (Figure 3B, Table 3). These results suggest that the interaction of the short hairpin loop of BrSCR9 with the hydrophobic cavity of BrSRK9 is important for activating SI, whereas the K206 residue is not essential for SI.

**Table 3.** SI phenotypes of *A. thaliana* transformants co-expressing different versions of *BrSRK-9 chimera* and *BrSCR-9* genes.

| Transgenes | Total no. of transformants | No. of self-pollen tubes per stigma<sup>a</sup> |  |  |
| --- | --- | --- | --- | --- |
|  |  | <10 | 10–29 | >29 |
| *BrSRK-9(K206L) chimera+BrSCR-9* | 16 | 10 | 0 | 6 |
| *BrSRK-9(V211E) chimera+BrSCR-9* | 19 | 0 | 0 | 19 |
| *BrSRK-9(P282V) chimera+BrSCR-9* | 13 | 6 | 1 | 6 |
| *BrSRK-9(P287I) chimera+BrSCR-9* | 15 | 0 | 0 | 15 |
| *BrSRK-9(F290H) chimera+BrSCR-9* | 17 | 0 | 0 | 17 |
| *BrSRK-9(P294M) chimera+BrSCR-9* | 20 | 0 | 0 | 20 |
| *BrSRK-9 chimera+BrSCR-9(K55N)* | 7 | 0 | 0 | 7 |
| *BrSRK-9 chimera+BrSCR-9(F69E)* | 13 | 0 | 0 | 13 |
| *BrSRK-9 chimera+BrSCR-9(Y73L)* | 8 | 0 | 0 | 8 |

<sup>a</sup>The number of pollen tubes per stigma was counted at 2 h after self-pollination. Values of <10, 10–29, and >29 indicate strong SI, weak SI, and compatible responses, respectively.

**Figure 3.**
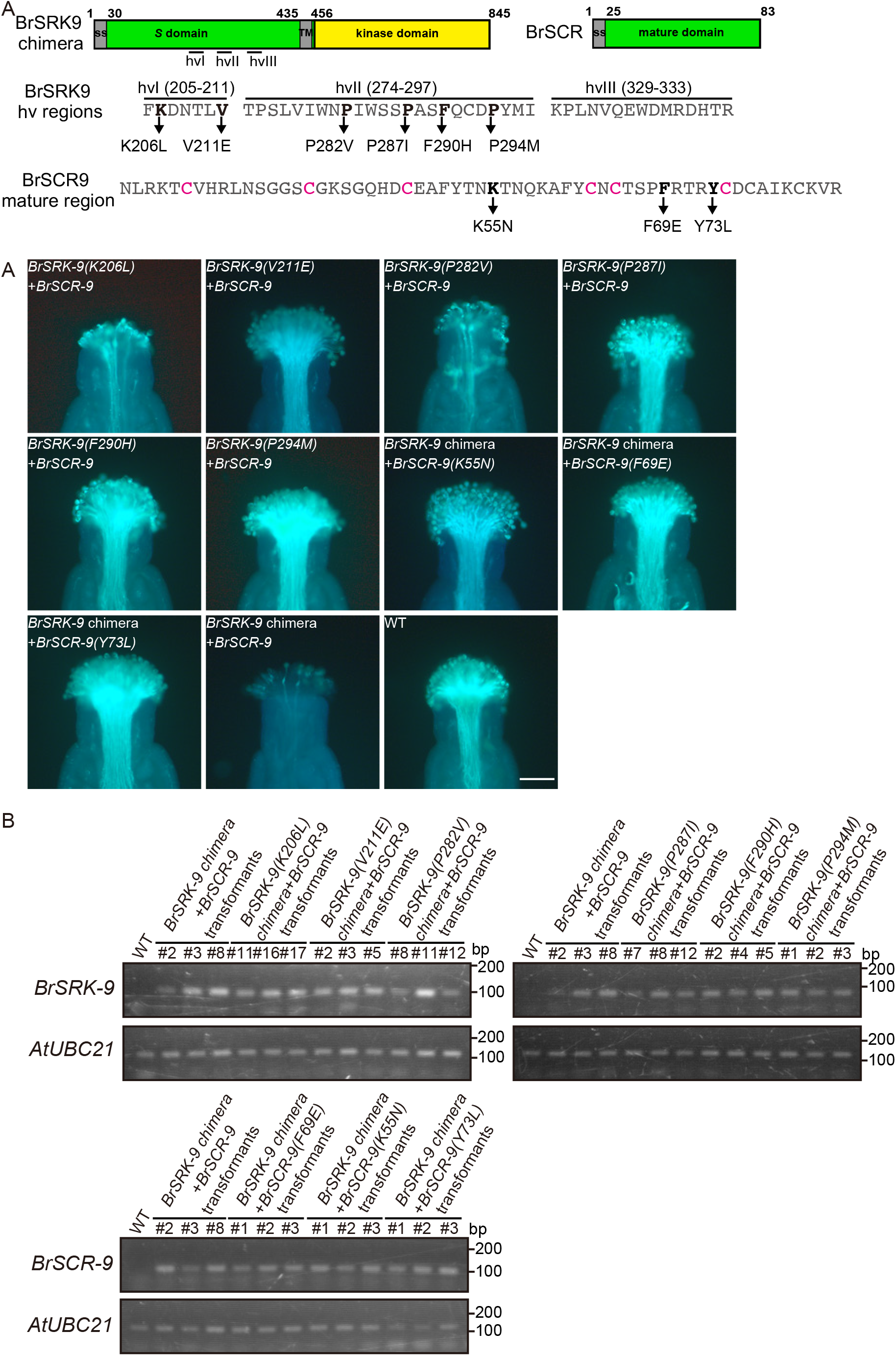
Functional analysis of mutant *BrSRK-9 chimera* and *BrSCR-9 in planta*. (A) Protein structure of BrSRK9 and BrSCR9 (top); amino acid sequences of hypervariable (hv) regions I (hvI), II (hvII), and III (hvIII) of BrSRK9 (middle); and amino acid sequence of the mature region of BrSCR9 (bottom). Letters in bold indicate amino acid residues used to introduce substitutions, and arrows indicate the amino acid substitutions. The conserved cysteine (C) residues of BrSCR are indicated in magenta. (B) Microscopic analysis of SI phenotypes. The genotype of *A. thaliana* plants used for pollination is indicated in each panel. Scale bar = 100 μm. (C) Expression analysis of *BrSRK-9 chimera* (upper panels) and *BrSCR-9* (lower panel) in wild-type and transgenic *A. thaliana* plants by reverse transcription PCR. The expression of *BrSRK-9 chimera* was examined in buds at positions −1 and −2, while that of *BrSCR-9* was examined in buds at positions −3 and −4. The plant genotype is indicated above each image. The *AtUBC21* gene was used as a control.

Next, we tested the importance of the amino acid residues of BrSRK9 and BrSCR9 located at the other eBrSRK9–BrSCR9 interaction sites. Lysine at position 55 (K55) of BrSCR9 located at the C-terminal end of α-helix has been reported to interact with the E325, D330, T332, and R333 of BrSRK9 [8]. We found that a mutation in the nucleotide sequence corresponding to the K55 residue of BrSCR9 abolished SI (Figure 3B, Table 3), suggesting that this interface is important for SI. Since it has been reported that the second lectin domain of BrSRK9 interacts with the α-helix of BrSCR9, and proline at position 282 (P282) of BrSRK9 located at the edge of this interface interacts with glycine at position 38 and arginine at position 70 of BrSCR9 [8], we constructed *A. thaliana* transformants expressing *BrSRK-9(P282V) chimera*+*BrSCR-9*. Six out of thirteen *BrSRK-9(P282V) chimera*+*BrSCR-9* transgenic *A. thaliana* lines showed SI (Figure 3B, Table 3), indicating that the P282V substitution in BrSRK9 does not cause a defect in SI.

To exclude the possibility that the nonfunctionality of the constructed mutants is due to no or low expression of mutant genes, we determined the expression of mutant variants of *BrSRK-9 chimera* and *BrSCR-9* by RT-PCR analysis. The results showed that transcript levels of the mutant *BrSRK-9 chimera* were as high as or higher than those of *BrSRK-9 chimera* and mutant (but functional) *BrSRK-9 chimera.* Similarly, transcript levels of mutant *BrSCR-9* were comparable to those of the wild-type *BrSCR-9* (Figure 3C).

## DISCUSSION

In the present study, we developed self-incompatible *A. thaliana* transformants showing the self-recognition activity of *B. rapa*. The majority of biochemical and structural findings related to self-recognition mechanisms in Brassicaceae have been obtained using *Brassica* species [5, 6, 8]. Our constructed *A. thaliana* transformants showing *B. rapa* self-recognition activity could be used to fill the gap between the results of *in vitro* analysis and those of *in planta* evaluations. We tested the importance of the amino acid residues required for the interaction between eBrSRK9 and BrSCR9 that have been predicted by structure analysis of the eBrSRK9–BrSCR9 complex [8] for activation of SI *in planta*. We found that some of the tested residues were essential (V211, P287, F290, and P294 of BrSRK9; and K55, F69, and Y73 of BrSCR9), whereas others were dispensable (K206 and P282 of BrSRK9) for SI. Interestingly, although K206 and P282 residues of BrSRK9 were not essential for SI, the K213 residue of CgSRK7 and AlSRK25 (which corresponds to the K206 residue of BrSRK9), and the M298 residue of AlSRK25 (which corresponds to the P282 residue of BrSRK9) (Supplemental Figure 5), are reportedly required for SI [16]. In addition, the result of F290H substitution in BrSRK9, which caused a defect in SI, was not consistent with the results of nucleotide mutations corresponding to the E292 and E295 residues of CgSRK7 and AlSRK25, respectively, both of which are not required for SI [16] (Supplemental Figure 5). These results suggest that the location of some of the amino acid residues of SRK, required for self-recognition activity, varies in the three-dimensional structure of SRK among the different *S* haplotypes. We speculate that this difference modifies the interface between SRK and SCR, and has contributed to the diversification of the recognition specificity of *S* haplotypes. Some of the *S* haplotypes in *B. rapa* are found to show high sequence similarity to those in *B. oleracea* and *Raphanus sativus*. Sequences of the *S* domain of SRK and mature region of SCR show 87.6% and 58.2% identity, respectively, between *B. rapa S-52* and *B. oleracea S-18* haplotypes, and both of these *S* haplotypes show distinct recognition specificities [6]. Comparison of *A. thaliana* transformants expressing *Brassica SRK chimera*+*SCR* with those expressing others with distinct recognition specificity and high sequence similarity, such as *BrSRK-52 chimera*+*BrSCR-52* and *BoSRK-18 chimera*+*BoSCR-18*, will help to analyze the mechanisms of differentiation of recognition specificity during the diversification of *S* haplotypes.

We found that *BrSRK-9 chimera*+*BrSCR-9* and *BrSRK-46 chimera*+*BrSCR-46* conferred SI in *A. thaliana*, whereas the remaining *BrSRK chimera*+*BrSCR* combinations belonging to *S-29*, *S-44*, and *S-60* haplotypes failed to confer SI. The *SRK* of *A. thaliana* ecotype Wei-1 has been reported to be nonfunctional in *A. thaliana* ecotype C24 due to the silencing of *SRK* by small RNAs produced from the inverted repeat structure present within the pseudo *SRK* gene in the C24 ecotype [37]. However, in this study, transcripts of *BrSRK-29*, *-44*, and *-60 chimera* accumulated to levels similar to those of the functional chimeric *SRK* transcripts, indicating that the cause of nonfunctionality of chimeric *B. rapa SRK* was not gene silencing. Our previous study revealed that AlSRK39, but not AlSRKb, functions in *A. thaliana* at high temperature, and this difference between AlSRK39 and AlSRKb was reflected by the different levels of accumulation of these plasma membrane-localized SRK proteins [34]. Therefore, we speculate that the nonfunctionality of chimeric *BrSRK-29*, *-44*, and *-60* in *A. thaliana* might be due to problems at the translational level, such as suboptimal accumulation of proteins, mislocalization of proteins, and failure of complex formation. Although we tried generating *A. thaliana* transformants expressing an epitope tag-fused BrSRK9 chimera along with BrSCR9, we were unable to obtain functional *A. thaliana* transformants showing SI, making it difficult to examine the subcellular localization of SRK. Therefore, construction of functional *A. thaliana* transformants expressing another epitope tag-fused BrSRK, and preparation of anti-BrSRK antibodies are required to examine the reason underlying the nonfunctionality of chimeric *BrSRK-29*, *-44*, and *-60* in *A. thaliana*.

In *B. rapa*, four class II *S* haplotypes (*S-29*, *S-40*, *S-44*, and *S-60*) have been reported [7]. We found that three of these four class II *S* haplotypes did not function in *A. thaliana*, suggesting that *Brassica* class II *S* haplotypes cannot confer SI in *A. thaliana*, although the possibility that other class II *S* haplotypes, such as *S-40*, function in *A. thaliana* cannot be excluded. Since more than 40 *S* haplotypes have been identified in *B. rapa* [31, 32], analysis of *A. thaliana* transformants expressing other *Brassica S* haplotypes will help identify the reason why different *Brassica S* haplotypes have different effects on SI in *A. thaliana*.

We found that the kinase domain of BrSRK is one of the main factors explaining why *BrSRK* is nonfunctional in *A. thaliana*. This result is consistent with the previous hypothesis that *Brassica* SI signaling differs from *Arabidopsis* SI signaling [14]. In *Brassica*, MLPK and ARC1 are reportedly phosphorylated by SRK [36, 38], suggesting that these two proteins are candidate substrates of *Brassica* SRK. On the other hand, substrate(s) of *Arabidopsis* SRK has not yet been reported. Identification of protein(s) phosphorylated by SRK in *Arabidopsis* could explain the differences in SI signaling between *Arabidopsis* and *Brassica*.

The present study suggests that the positions of amino acid residues of SRK required for self-recognition activity vary among the different *S* haplotypes. We also found that the ability of a *Brassica S* haplotype to confer SI in *A. thaliana* differs from that of the other *S* haplotypes. In addition, we previously reported that different *S* haplotypes show different responses to high temperature [34]. These findings suggest that analysis of multiple *S* haplotypes is important for understanding the mechanisms of SI. Therefore, the method developed in this study to construct self-incompatible *A. thaliana* plants showing *Brassica* self-recognition activity contributes to increasing the number of *S* haplotypes that can be analyzed in *A. thaliana*, which will help to elucidate the molecular mechanisms of SI.

## Supporting information

Supplememtal information, and will be used for the link to the file on the preprint site.

## ACKNOWLEDGMENTS

We thank Kai Miyakoshi for cloning of *BrSCR-9*, *BrSRK-29*, *BrSCR-29*, *BrSRK-44*, *BrSCR-44*, *BrSRK-46*, and *BrSCR-46*. We also thank members of the Plant Breeding and Genetics laboratory in Tohoku University for constructing the *B. rapa S*-tester lines. This work was supported in part by JSPS KAKENHI (grant numbers 26892004 and 20K05979 to M.Y.).

## AUTHOR CONTRIBUTIONS

M.Y. designed and preformed the experiments. M.Y., H.K., and T.N. analyzed the data and wrote the manuscript.

## MATERIALS and METHODS

### Plant materials and growth conditions

Plants of *Arabidopsis thaliana* ecotype C24 obtained from the Arabidopsis Biological Resource Center (ABRC; https://abrc.osu.edu/) were used as the wild type (WT). *AlSRK-b*+*AlSCR-b A. thaliana* transformants have been described previously [34]. Seeds were sterilized with 20% bleach and sown on Murashige and Skoog (MS) medium (Wako, Osaka, Japan) containing 0.7% (w/v) agar and 1% (w/v) sucrose. All *A. thaliana* plants were grown at a constant temperature of 23°C under long-day photoperiod (16 h light/8 h dark) and 100 μmol photons m^−2^ s^−1^ light intensity.

### Plasmid construction and plant transformation

The *pMYC192* plasmid (containing the *AlSCR-b* promoter and *AlSCR-b* terminator) and *pMYC290* plasmid (containing the stigma papilla cell-specific promoter of *A. thaliana* [*AtS1*; *At3g12000*] and *AlSRK-b* terminator) have been described previously [34].

The *BrSRK-9* coding sequence (CDS) fragments were amplified by PCR using first-strand cDNA (template; synthesized from total RNA isolated from the stigma of the *B. rapa S-9* haplotype) and the BrSRK9-F/BrSRK9-R primer pair. The *BrSRK-9* CDS fragment was introduced into the *Kpn*I site of *pMYC290* by InFusion cloning (TAKARA Bio, Shiga, Japan). Fragments of CDS of the *BrSRK S* domain and transmembrane domain were amplified by PCR using the first-strand cDNA synthesized from stigma RNA (template) with gene-specific primers (BrSRK9-F/BrSRK9(S)-R for *BrS-9*; BrSRK29-F/BrSRK29_44(S)-R for *BrS-29*; BrSRK44-F/BrSRK29_44(S)-R for *BrS-44*; BrSRK46-F/BrSRK46(S)-R for *BrS-46*; and BrSRK60-F/BrSRK60(S)-R for *BrS-60*) (Supplemental Table 4). Fragments of *AlSRK-b* kinase domain CDS were amplified from the entire *AlSRK-b* CDS by PCR using sequence-specific primers (AlSRKb(kin)-F9/AlSRKb-R for *BrS-9*; AlSRKb(kin)-F29_44/AlSRKb-R for *BrS-29* and *BrS-44*; AlSRKb(kin)-F46/AlSRKb-R for *BrS-46*; and AlSRKb(kin)-F60/AlSRKb-R for *BrS-60*). The amplified DNA fragments were pooled and used as a template to synthesize *BrSRK chimera* fragments by PCR using sequence-specific primers (BrSRK9-F/AlSRKb-R for *BrS-9*; BrSRK29-F/AlSRKb-R for *BrS-29*; BrSRK44-F/AlSRKb-R for *BrS-44*; BrSRK46-F/AlSRKb-R for *BrS-46*; and BrSRK60-F/AlSRKb-R for *BrS-60*). The amplified *BrSRK-9* and *BrSRK chimera* fragments were introduced into the *Kpn*I site of *pMYC290* by InFusion cloning (TAKARA Bio).

To express *BrSCR* in *A. thaliana* anthers, a DNA cassette containing the *AlSCR-b* promoter, *BrSCR* CDS, and *AlSCR-b* terminator was synthesized. Briefly, fragments of the *BrSCR* CDS were amplified by PCR using first-strand cDNA (template; synthesized from the total RNA of *B. rapa* anthers) and gene-specific primers (BrSCR9-F/BrSCR9-R for *BrS-9*; BrSCR46-F/BrSCR46-R for *BrS-46*; and BrSCRII-F/BrSCRII-R for *BrS-29*, *BrS-44*, and *BrS-60*) (Supplemental Table 4). The *BrSCR-9* CDS fragment was introduced into the *Kpn*I/*Sal*I sites of *pMYC192*, and *BrSCR-29*, *-44*, *-46*, and *-60* CDS fragments were introduced into the *Kpn*I/*Pst*I sites of *pMYC192*. The constructed plasmids were used as templates for PCR using sequence-specific primer sets: AlSCRbpro(H)-F/AlSCRbterm(H)-R for *BrSCR-9* and -*46*, and AlSCRbpro(P)-F/AlSCRbterm(P)-R for *BrSCR-29*, -*44*, and -*60*. The amplified *BrSCR-9* cassette was introduced into the *Hin*dIII site of the constructed plasmid containing the *BrSRK-9* and *BrSRK-9 chimera*, and the *BrSCR*-*46* cassette was similarly cloned into the plasmid containing the *BrSRK-46 chimera* by InFusion cloning (TAKARA Bio). Similarly, the *BrSCR-29*, *-44*, and *-60* cassette was introduced into the *Pst*I site of the constructed plasmid containing the *BrSRK-29*, *-44*, and *-60 chimera*, respectively, by InFusion cloning (TAKARA Bio). To introduce mutations in the *BrSRK-9 chimera* and *BrSCR-9* sequences, the plasmid containing *BrSRK-9 chimera+BrSCR-9* genes was used as a template for recombinant PCR-mediated site-directed mutagenesis with primers listed in Supplemental Table 4.

To express the *BrMLPK* gene in *A. thaliana*, the *AtS1* promoter fragments were amplified by PCR from the *AlSRK-b:FLAG*+*AlSCR-b* plasmid [39] DNA (template) using the AtS1-F/AtS1-R primer pair. The amplified fragment was introduced into the *Pst*I/*Xba*I sites of *pRI201-AN* (TAKARA Bio), and the construct was designated as *pMYC241*. The *Nde*I site of *pMYC241* was replaced with the *Stu*I site by PCR. In the first-round PCR, *pMYC241* was used as a template, and AtS1-F/pRI201-StuI_R and pRI201-StuI_F/M13_R primer pairs were used to amplify the first and second fragments, respectively. Then, the first-round PCR products were used as templates with the AtS1-F/M13_R primer pair. The resulting PCR product was introduced into the *Xba*I/*Sac*I sites of *pMYC241*, and the construct was designated as *pMYC242*. The *BrMLPK* gene fragments were amplified by PCR using genome DNA extracted from the leaves of *B. rapa S-9* haplotype by the cetyl trimethyl ammmonium bromide (CTAB) method [40] as a template and the MLPK_InFusionF/MLPK_InFusionR primer pair. The amplified fragment was introduced into the *Stu*I site of *pMYC242* by InFusion cloning (TAKARA Bio), and the construct was designated as *pMYC266*.

To verify the absence of PCR-generated polymorphisms, all constructed plasmids were sequenced on the Beckman Coulter CEQ2000XL DNA sequencer (SCIEX, Framingham, MA) using the GenomeLab DTCS Quick Start Kit (SCIEX). The sequence-verified plasmids were introduced into *Agrobacterium tumefaciens* strain GV3101 [41], which was then used to transform *A. thaliana* plants using the floral dip method [42]; the *AtS1*_*pro*_:*BrMLPK* construct was introduced into the *BrSRK-9*+*BrSCR-9* line #1, whereas the remaining constructs were introduced into WT C24 plants.

### Pollination assays and seed counting

In the pollination assay, flowers were emasculated just before flowering, which corresponds to the flowering stage 13 in *A. thaliana* [43]. The emasculated flowers were collected and placed on a 0.5% agar plate. Stigmas were manually pollinated under a stereomicroscope with pollen grains from open flowers. The pollinated stigmas were incubated at 23°C under continuous light for 2 h and then fixed in an ethanol:acetic acid (3:1) solution for 10 min at 55°C. The fixed stigmas were treated with 8 M NaOH at room temperature for 15 min and then stained with decolorized aniline blue. The stained stigmas were observed by fluorescence microscopy, as previously described [44]. Each pollination assay was performed using at least three stigmas. The degree of SI activity was classified according to the number of pollen tubes per pollinated stigma as follows: strong SI, <10; weak SI, 10–29; compatible, >29. Images of pollinated stigmas were captured using an Axioskop microscope fitted with an AxioCam ERc 5s camera (Carl Zeiss, Oberkochen, Germany).

To count the number of seeds per silique, 10 siliques between positions 6 and 15 were collected from the main stem. The number of seeds per silique was counted under a stereomicroscope. This experiment was performed using three plants per transgenic line.

### Gene expression analysis

Total RNA was isolated using the TRIzol reagent (Thermo Fisher Scientific, Waltham, MA) from 10 *A. thaliana* flowers immediately after flowering and at the flowering stage (+1 and 0 positions, respectively), and from buds immediately before flowering and the second bud before flowering (−1 and −2 positions, respectively), the third and fourth buds before flowering (−3, −4 positions, respectively), and the fifth and sixth buds before flowering (−5, −6 positions, respectively). The isolated total RNA was treated with RQ1 RNase-free DNase (Promega, Fitchburg, WI), and first-strand cDNA was synthesized using the PrimeScript II 1^st^ strand cDNA Synthesis Kit (TAKARA Bio), according to the manufacturer’s protocol.

RT-PCR was performed using homemade *Taq* [45] with gene-specific primers (Supplemental Table 5) under the following amplification conditions: 94°C for 2 min, followed by 30 cycles of 94°C for 30 s, 58°C for 30 s, and 72°C for 30 s. qRT-PCR was performed on the CFX Connect Real-Time System (Bio-Rad) using the THUNDERBIRD SYBR qPCR mix (TOYOBO, Osaka, Japan) with 40 cycles of denaturation at 95°C for 15 s and extension at 60°C for 60 s. The transcript abundances of *AlSRK-b* and *AlSCR-b* were compared at different flower/bud developmental stages using the ΔΔC method. The transcript amounts of *AlSRK-b*, *BrSRK-9*, *BrSRK-9 chimera*, *BrSRK-29 chimera*, *BrSRK-44chimera*, *BrSRK-46 chimera*, *BrSRK-60 chimera*, *AlSCR-b*, *BrSCR-9*, *BrSCR-29*, *BrSCR-44*, *BrSCR-46*, and *BrSCR-60* were determined using the standard curve method. Standard double-stranded DNAs for the standard curve method were produced from the synthesized first-strand cDNA by PCR using gene-specific primers (Supplemental Table 5). *AtUBC21* (*At5g25760*) was used as an internal control to normalize the RNA quantity in both ΔΔCT and standard curve methods. Each sample was analyzed in three biological replicates, each containing three technical repeats.

### Sequence analysis

SignalP-5.0 [46] and SOSUIsignal [47] were used to predict the signal sequences of SRK and SCR, and the transmembrane regions of SRK, respectively. Sequences were aligned using Clustal Omega [48].

## Accession numbers

Sequence data from this study can be found in the GenBank/EMBL data libraries under the following accession numbers: AB052756 (*AlSRK-b*), AB052754 (*AlSCR-b*), LC556298 (*BrSRK-9*), AB022078 (*BrSCR-9*), AB008191 (*BrSRK-29*), AB067449 (*BrSCR-29*), NEW (*BrSRK-44*), LC556299 (*BrSCR-44*), AB257128 (*BrSRK-46* and *BrSCR-46*), AB097116 (*BrSRK-60*), AB067446 (*BrSCR-60*), EF530735 (*CgSRK-7*), GQ351355 (*AlSRK-25*), and AC189432/AB121973/EF179103 (*BrMLPK*).

## SUPPLEMENTAL INFORMATION

**Supplemental Figure 1.** Quantitative real-time PCR (qRT-PCR) analysis of *AlSRK-b*+*AlSCR-b* transgenic *Arabidopsis thaliana* floral buds at different positions. Expression levels of *AlSRK-b* (upper graphs; blue) and *AlSCR-b* (lower graphs; magenta) in *AlSRK-b*+*AlSCR-b* transgenic *A. thaliana* buds at positions +1 and 0 [+1, 0], −1 and −2 [−1, −2], −3 and −4 [−3, −4], and −5 and −6 [−5, −6] were compared among flower/bud developmental stages using the ΔΔCT method, and the relative expression level of each gene in buds at positions −1 and −2 was set at 1. *AtUBC21* was used as an internal control. Data represent mean ± standard deviation (SD) of three biological replicates. Different letters indicate significant differences (*P* < 0.05; Tukey–Kramer method). Expression data are listed in Supplemental Table 1.

**Supplemental Figure 2.** Absolute qRT-PCR analysis of transgenic *A. thaliana* lines expressing *Arabidopsis lyrata S-b* and *Brassica rapa S* genes.

Expression levels of *SRK* (upper graph; blue) and *SCR* (lower graph; magenta) were analyzed by absolute qRT-PCR in various *A. thaliana* transformants (indicated below the graph) in buds at positions −1 and −2 and at positions −3 and −4, respectively. The *AtUBC21* transcript level was set at 1 in each experiment. Data represent mean ± SD of three biological replicates. Different letters indicate significant differences (*P* < 0.05; Tukey–Kramer method). Expression data of *SRK* and *SCR* are listed in Supplemental Table 2 and Supplemental Table 3, respectively.

**Supplemental Figure 3.** Amino acid sequence alignment of AlSRKb and BrSRKs.

Amino acid sequences of AlSRKb, BrSRK9, BrSRK46, BrSRK29, BrSRK44, and BrSRK60 were aligned using Clustal Omega. Signal sequences predicted using SignalP-5.0 are shown in green, and transmembrane regions predicted using SOSUIsignal are shown in red. Hypervariable (hv) regions I (hvI)), II (hvII), and III (hvIII) are outlined in red boxes. Amino acid residues of BrSRK9 mutants analyzed in Figure 4 are indicated in cyan. The kinase domain of BrSRK replaced by that of AlSRKb in BrSRK chimeras is indicated by a purple line.

**Supplemental Figure 4.** *BrMLPK* does not affect the self-incompatibility (SI) response of *BrSRK-9*+*BrSCR-9* transgenic *A. thaliana* plants.

(A) Self-pollination assay of wild-type (WT) and transgenic *A. thaliana* plants expressing *BrSRK-9*, *BrSCR-9*, and *BrMLPK* (*BrSRK-9*+*BrSCR-9*+*BrMLPK*). A large number of self-pollen tubes were observed in the stigma of transgenic plants, similar to that observed in the stigma of wild-type plants. Scale bar = 100 μm.

(B) RT-PCR analysis of *BrSRK-9* and *BrMLPK* in the buds (positions −1 and −2) of wild-type (WT) plants, *BrSRK-9*+*BrSCR-9* transgenic *A. thaliana* line #1 (*BrSRK-9*+*BrSCR-9* #1), and *BrSRK-9*+*BrSCR-9*+*BrMLPK A. thaliana* lines #1, #2, and #3 (*BrSRK-9*+*BrSCR-9*+*BrMLPK* #1, #2, and #3). *AtUBC21* was used as an internal control.

**Supplemental Figure 5.** Amino acid sequence alignment of BrSRK9, *Capsella grandiflora* SRK7, and AlSRK25.

Amino acid sequences flanking the hv regions (outlined in red boxes) of BrSRK9, CgSRK7, and AlSRK25 were aligned using Clustal Omega. Amino acid residues essential and non-essential for SI are indicated in magenta and green, respectively.

**Supplemental Table 1.** qRT-PCR analysis of *Arabidopsis lyrata S-b* genes in transgenic *Arabidopsis thaliana* floral buds at different positions.

**Supplemental Table 2.** Absolute qRT-PCR analysis of transgenic *A. thaliana* plants expressing *AlSRK* and *Brassica rapa SRK* genes. Related to Tables 1 and 2 and Figures 1 and 2

**Supplemental Table 3.** Absolute qRT-PCR analysis of transgenic *A. thaliana* plants expressing *AlSCR* and *BrSCR* genes.

**Supplemental Table 4.** List of primers used for plasmid construction.

**Supplemental Table 5.** List of primers used for absolute qRT-PCR and reverse-transcription PCR (RT-PCR).

