## Supplementary material for "Development of *Arabidopsis thaliana* transformants showing the self-recognition activity of *Brassica rapa*": Supplememtal information, and will be used for the link to the file on the preprint site.

Supplemental Figure 1 Yamamoto et al.

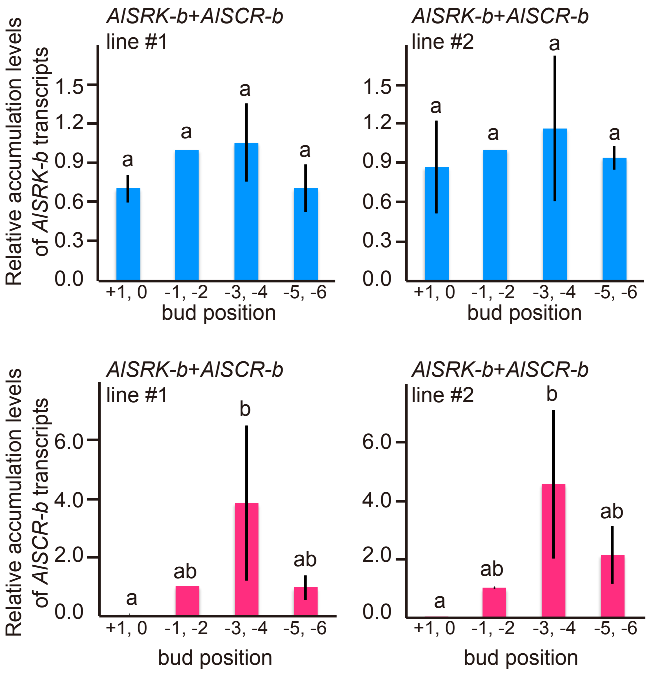


**Supplemental Figure 1.** Quantitative real-time PCR (qRT-PCR) analysis of *AlSRK-b+AlSCR-b* transgenic *Arabidopsis thaliana* floral buds at different positions. Expression levels of *AlSRK-b* (upper graphs; blue) and *AlSCR-b* (lower graphs; magenta) in *AlSRK-b+AlSCR-b* transgenic *A. thaliana* buds at positions +1 and 0 [+1, 0], -1 and -2 [-1, -2], -3 and -4 [-3, -4], and -5 and -6 [-5, -6] were compared among flower/bud developmental stages using the ΔΔCT method, and the relative expression level of each gene in buds at positions -1 and -2 was set at 1. *AtUBC21* was used as an internal control. Data represent mean ± standard deviation (SD) of three biological replicates. Different letters indicate significant differences ($P < 0.05$; Tukey–Kramer method). Expression data are listed in Supplemental Table 1.

Supplemental Figure 2 Yamamoto et al.

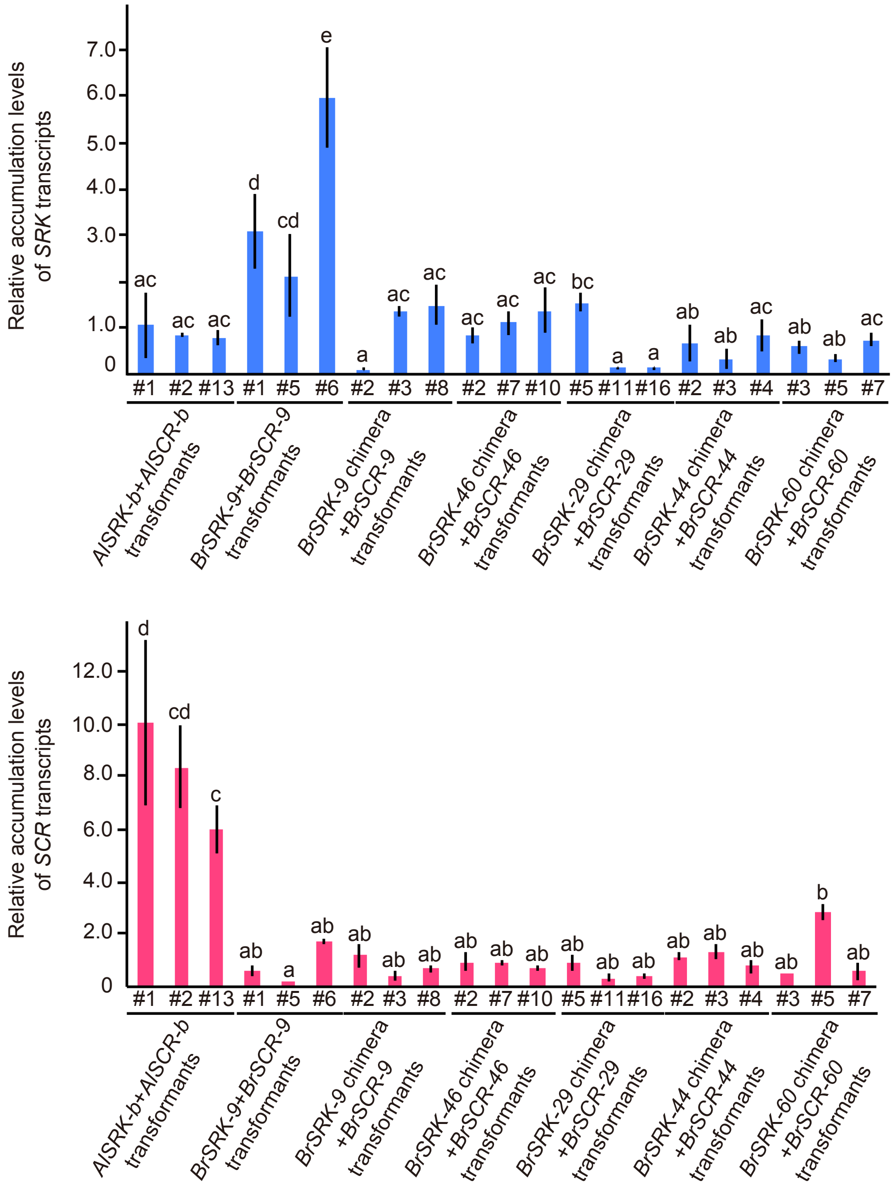


**Supplemental Figure 2.** Absolute qRT-PCR analysis of transgenic *A. thaliana* lines expressing *Arabidopsis lyrata S-b* and *Brassica rapa S* genes.
Expression levels of *SRK* (upper graph; blue) and *SCR* (lower graph; magenta) were analyzed by absolute qRT-PCR in various *A. thaliana* transformants (indicated below the graph) in buds at positions -1 and -2 and at positions -3 and -4, respectively. The *AtUBC21* transcript level was set at 1 in each experiment. Data represent mean ± SD of three biological replicates. Different letters indicate significant differences ($P < 0.05$; Tukey–Kramer method). Expression data of *SRK* and *SCR* are listed in Supplemental Table 2 and Supplemental Table 3, respectively.

```
AlSRKb        MRVVVPNCHH-----FYIFFVVLILIRSVFSSYVHTLSSTESLTISSKQTIVSPGEVFEL    55
BrSRK9        MKGVRNIYHHSYT--FLLVFFVMILFRPAFSL--STLSSTESLTISSNRTLVSPGNIFEL    56
BrSRK46       MKGVRNIYHHSYT-SFLLVFVVMILFRPTLSIYFNTLSSTESLTISNSRTLVSPGDVFEL    59
BrSRK29       MKRVQNIYHHSYT-SFLLVFLVLILFHPALSIYVNTMSSSESLTISSNRTLVSPGGVFEL    59
BrSRK44       MKRVKNIYHH-YTFSFLLVFLVLILFHPALSIYVNTMSSSESLTISSNRTLVSPGGVFEL    59
BrSRK60       MKGVHNIYHHSYTFSFLLVFLALILFHPALSTYVNTMSSSESLTISSNRTLVSPGGVFEL    60
              *: *    **     * :.*..:**:: .:*    *:**:******..:*:**** :***

AlSRKb        GFFNPAATSRDGDRWYLGIWFKTNLERTYVWVANRDNPLYNSTGTLKISDTNLVLLDQFD    115
BrSRK9        GFFRT--N----SRWYLGMWYKKLSGRTYVWVANRDNPLSNSIGTLKISNMNLVLLDHSN    110
BrSRK46       GFFKTTSS----SRWYLGIWYKKLPGRTYVWVANRDNPLSNSIGTLKISNMNLVILDHSN    115
BrSRK29       GFFKTLER----SRWYLGIWYKKVPWKTYAWVANRDNPLSNSIGTLKISGNNLVLLGQSN    115
BrSRK44       GFFKPLGR----SRWYLGIWYIKVPLKTYAWVANRDNPLSSSIGTLKISGNNLVLLGQSN    115
BrSRK60       GFFKPSGR----SRWYLGIWYKKVSQKTYAWVANRDNPLSNSIGTLKISGNNLVLLGQSN    116
              ***.        .*****:*: .   :**.********* .* ******. ***:*.: :

AlSRKb        TLVWSTNLTG-VLRSPVVAELLSNGNLVLKDSKTNDKDGILWQSFDYPTDTLLPQMKMGW    174
BrSRK9        KSVWSTNLTRENVRSPVVAELLANGNFVVR-----DPSGFLWQSFDYPTDTLLPEMKLGY    165
BrSRK46       KSVWSTNHTRGNERSLVVAELLANGNFLMRDSNSNDAYGFLWQSFDYPTDTLLPEMKLGY    175
BrSRK29       NTVWSTNFTRGNARSPVIAELLPNGNFVMRHSNNKDSNGFLWQSFDFPTDTLLPEMKLGY    175
BrSRK44       NTVWSTNLTRGNARSPVIAELLPNGNFVIRHSNNKDSSGFLWQSFDFPTDTLLPEMKLGY    175
BrSRK60       NTVWSTNLTRENVRSPVIAELLPNGNFVMRYSNNKDSSGFLWQSFDFPTDTLLPEMKLGY    176
              . ***** *    ** *:**** ***::::     *  *:******:*******:**:*:

AlSRKb        DVKKGLNRFLRSWKSQYDPSSGDFSYKLET-RGFPEFFLLWR----NSRVFRSGPWDGLR    229
BrSRK9        DLKTGLNRFLVSWRSSDDPSSGDFSYKLDIQRGLPEFYTFKD----NTLVHRTGPWNGIR    221
BrSRK46       DLKIGLNRSLTSWRSPDDPSSGYFSYKLEGSRRLPEFYLMQG----DVREHRSGPWNGIQ    231
BrSRK29       NLKTGRNRFLTSWKSSDDPSSGNFAYKLDLRRGLPEFILINTFLNQRVETQRSGPWNGME    235
BrSRK44       DLKTGRNRFLTSWKGSDDPSRGNFVYKLDIRRGLPEFILINQFLNQRVETQRSGPWNGME    235
BrSRK60       DFKTGRNRFLTSWRSYDDPSSGKFTYELDIQTGLPEFILINRFLNQRVVMQRSGPWNGIE    236
              :.* * ** * **:.  *** * * *:*:    :***  :           *:***:*:.
                                                   hvI

AlSRKb        FSGIPEMQQWEYMVSNFTENREEVAYTFQITNHNIYSRFTMSSTGALKRFRWISSSEEWN    289
BrSRK9        FSGIPEEQQLSYMVYNFTENSEEVAYTFLVTNNSIYSRLTINFSGFFERLTWTPSLVIWN    281
BrSRK46       FIGIPEDQKSSYMMYNFTDNSEEVAYTFVMTNNGIYSRLKLSSDGYLERLTWAPSSGAWN    291
BrSRK29       FSGIPEVQGLNYMVYNYTENSEEISYSFHMTNQSIYSRLTVS-ELTLNRFTWIPPSSAWS    294
BrSRK44       FSGIPEVQGLNYMVYNYTENSEEISYSFHMTNQSIYSRLTVS-EFTFDRLTWIPPSRDWS    294
BrSRK60       FSGIPEVQGLNYMVYNYTENSEEIAYSFQMTNQSIYSRLTVS-DYTLNRFTRIPPSWGWS    295
              * **** *  .**: *:*:* **::*:* :**:.****:.:.    :.*:        *.

AlSRKb        QLWNKPN-DHCDMYKRCGPYSYCDMNTSPICNCIGGFKPRNLHEWTLRNGSIGCVRKTRL    348
BrSRK9        PIWSSPASFQCDPYMICGPGSYCDVNTLPLCNCIQGFKPLNVQEWDMRDHTRGCIRRTRL    341
BrSRK46       VFWSSPN-HQCDMYRMCGTYSYCDVNTSPSCNCIPGFNPKNRQQWDLRIPISGCKRRTRL    350
BrSRK29       LFWTLPT-DVCDPLYLCGSYSYCDLITSPNCNCIRGFVPKNPQQWDLRDGTQGCVRTTQM    353
BrSRK44       LFWTLPT-DVCDPLYLCGSYSYCDLITSPNCNCIRGFVPKNPQQWDLRDGTQGCVRRTQM    353
BrSRK60       LFWSLPT-DVCDSLYFCGSYSYCDLNTSPYCNCIRGFVPKNRQRWDLRDGSHGCVRTTQM    354
               :*. *    **    **  ****: * * **** ** * * :.* :*    ** * *::
                     hvII                                      hvIII

AlSRKb        NCGGDGFLCLRKMKLPDSLAAIVDRTIDLGECKKRCLNDCNCTAYASTDIQNGGLGCVIW    408
BrSRK9        SCRGDGFTRMKNMKLPETTMATVDRSIGVKECEKKCLSDCNCTAFANADIRDGGTGCVIW    401
BrSRK46       SCNGDGFTRMKNMKLPDTTMAIVDRSMGVKECEKRCLSDCNCTAFANADIRNGGTGCVIW    410
BrSRK29       SCSGDGFLRLNNMNLPDTKTATVDRTIDVKKCEERCLSDCNCTSFAAADVRNGGLGCVFW    413
BrSRK44       SCGRDGFLRLNNMNLPDTKTATVDRTMDVKKCEERCLSDCNCTSFAAADVKNGGIGCVFW    413
BrSRK60       SCSGDGFLRLNNMNLPDTKTASVDRTIDVKKCEEKCLSDCNCTSFATADVRNGGLGCVFW    414
              .*  ***  :.:*:**::  * ***::.: :*:::**.*****::* :*:::** ***:*
```

**Supplemental Figure 3.** Amino acid sequence alignment of AlSRKb and BrSRKs.
Amino acid sequences of AlSRKb, BrSRK9, BrSRK46, BrSRK29, BrSRK44, and BrSRK60 were aligned using Clustal Omega. Signal sequences predicted using SignalP-5.0 are shown in green, and transmembrane regions predicted using SOSUIsignal are shown in red. Hypervariable (hv) regions I (hvI)), II (hvII), and III (hvIII) are outlined in red boxes. Amino acid residues of BrSRK9 mutants analyzed in Figure 4 are indicated in cyan. The kinase domain of BrSRK replaced by that of AlSRKb in BrSRK chimeras is indicated by a purple line.

Supplemental Figure 3 Yamamoto et al.

```
AlSRKb          IEELLDIRNYASGGQDLYVRLADVD----IGDERNIRGKIIGLAVGASVILF---LSSIM     461
BrSRK9          TGRLDDMRNYAVSGQDLYVRLAAAD----VVEKRTANGKIVSLIVGVCVL-----LLLIF     452
BrSRK46         TGELEDMRNYAEGGQELYVRLAAAD----LVKKRNGNWKIISLIVGVSVVLLLLLLLLIM     466
BrSRK29         TGELVAIRKFAVGGQDLYVRLNAADLDLSSGEKRDRTGKIIGWSIGVSVMLI---LSVIV     470
BrSRK44         TGELVAIRKFAVGGQDLYVRLNAADLDISSGEKRDRTGKIIGWSIGVSVMLI---LSVIV     470
BrSRK60         TGDLVEIRKQAVVGQDLYVRLNAADLDFSSGEKRDRTGTIIGWSIGVSVMLI---LSVIV     471
                   *  :*: *  **:*****  .*      .:*    .*:.  :*..*:     *  *.

AlSRKb          FCVWRRKQKLLRATEAPIVYPTINQGLLMNRLEI-SSGRHLSEDNQTEDLELPLVEFEAV     520
BrSRK9          FCLWKRKQRRAKAMATSIVHRQRKQILLMNGMTL-SNNRQLSRENKTGEFELPLIELEAV     511
BrSRK46         FCLWKRKQNRAKAMATSIVNQQRNQNVLMNTMTQ-SNKRQLSRENKADEFELPLIELEAV     525
BrSRK29         FCFWRRKHKQAKADATPIVGNQ----VLMNEVVLPRKKRNFSGEDEVENLELPLMEFEAV     526
BrSRK44         FCFWRRRQKQAKADATPIVGNQ----VLMNEVVLPRKKRNFSGEDEVENLELPLMEFEAV     526
BrSRK60         FCFWRRRQKQAKADATPIVGNQ----VLMNEVVLPRKKIHFSGEDEVENLELSLMEFEAV     527
                **.*:*::.  :*  : **       :*** :    .  ::* :::. ::** *:*:***

AlSRKb          VMATENFSNSNKLGEGGFGVVYKGRLLDGQEIAVKRLSTTSIQGICEFRNEVKLISKLQH     580
BrSRK9          VKSTENFSNCNKLGQGGFGIVYKGT-LDGQEIAVKRLSKTSVQGADEFMNEVTLIARLQH     570
BrSRK46         VKATENFSNCNELGRGGFGIVYKGM-LDGQEVAVKRLSKTSLQGIDEFMNEVRLIARLQH     584
BrSRK29         VTATEHFSDFNKVGKGGFGVVYKGRLVDGQEIAVKRLSEMSAQGTDEFMNEVRLIAKLQH     586
BrSRK44         VTATEHFSDLNKVGKGGFGVVYKGRLVDGQEIAVKRLSEMSAQGTDEFMNEVRLIAKLQH     586
BrSRK60         VTATEHFSDFNKVGKGGFGVVYKGRLVDGQEIAVKRLSEMSAQGTDEFMNEVRLIAKLQH     587
                * :**:**: *::*.****:****  :****:******  * **  ** *** **::***

AlSRKb          INLVRLFGCCVDENEKMLIYEYLENLSLDSHLFNKSLSCKLNWQMRFDITNGIARGLLYL     640
BrSRK9          INLVQILGCCIDADEKMLIYEYLENLSLDSYLFGKTRSSKLNWKERFDITNGIARGLLYL     630
BrSRK46         INLVRILGCCIEAGEKILIYEYLENSSLDYFLFGKKRSSNLNWKDRFAITNGVARGLLYL     644
BrSRK29         NNLVRLLGCCVYEGEKILIYEYLENLSLDSHLFDGSRSCKLNWQMRFDIINGIARGLLYL     646
BrSRK44         NNLVRLLGCCVYEGEKILIYEYLENLSLDSHLFDETRSCMLNWQMRFDIISGIARGLLYL     646
BrSRK60         NNLVRLLGCCVYEGEKILIYEYLENLSLDSHLFDETRSCMLNWQMRFDIINGIARGLLYL     647
                 ***:::***:  .**:******** *** .**. . *. ***: ** * .*:*******

AlSRKb          HQDSRFRIIHRDLKASNVLLDKDMTPKISDFGMARIFGRDETEANTRKVVGTYGYMSPEY     700
BrSRK9          HQDSRFRIIHRDLKVSNILLDKNMIPKISDFGMARIFARDETEANTMRVVGTYGYMSPEY     690
BrSRK46         HQDSRFRIIHRDLKPGNILLDKYMIPKISDFGMARIFARDETQVRTDNAVGTYGYMSPEY     704
BrSRK29         HQDSRFRIIHRDLKASNVLLDKDMTPKISDFGMARIFGRDETEADTRKVVGTYGYMSPEY     706
BrSRK44         HQDSRFRIIHRDLKASNVLLDKDMTPKISDFGMARIFGRDETEADTRKVVGTYGYMSPEY     706
BrSRK60         HQDSRFRIIHRDLKASNVLLDKDMTPKISDFGMARIFGQDETEADTRKVVGTYGYMSPEY     707
                ************** .*:**** * ************.:***:. * ..***********

AlSRKb          AMDGIFSVKSDVFSFGVLVLEIVSGKKNRGFYNSNQDNNLLGYAWRNWKEGKGLEILDPF     760
BrSRK9          AMEGIFSEKSDVFSFGVIVLEIVTGKRNREF---NNENNLLSYAWSNWKEGRALEIVDPD     747
BrSRK46         AMYGVISEKTDVFSFGVIVLEIVIGKRNRGFYQVNPENNLPSYAWTHWAEGRALEIVDPV     764
BrSRK29         AMNGTFSMKSDVFSFGVLLLEIISGKRNKGFCDSDSSLNLLGCVWRNWKEGQGLEIVDRV     766
BrSRK44         AMNGTFSMKSDVFSFGVLLLEIISGKRNKGFCDSDSTLNLLGCVWRNWKEGQGLEIVDKF     766
BrSRK60         AMNGTFSMKSDVFSFGVLLLEIISGKRNKGFCDSDSNLNLLGCVWRNWKEGQGLEIVDRV     767
                ** * :* *:*******::***: **:*: *   :   ** . .* :* **:.***:*

AlSRKb          IVDSSS-SPSAFRPHEVLRCIQIGLLCVQERAEDRPVMSSVVVMLRSETETIPQPKPPGY     819
BrSRK9          IVDSLSPLSSTFQPQEVLKCIQIGLLCVQELAEHRPTMSSVVWMLGSEATEIPQPKPPGY     807
BrSRK46         ILDSLSSLPSTFKPKEVLKCIQIGLLCIQERAEHRPTMSSVVWMLGSEATEIPQPKPPVY     824
BrSRK29         IIDSSS--P-TFRPSEISRCLQIGLLCVQERVEDRPMMSSVVLMLGSEAALIPQPKQPGY     823
BrSRK44         INDSSS--P-TFKPREILRCLQIGLLCVQERVEDRPMMSSVVLMLGSEAALIPQPKQPGY     823
BrSRK60         IIDSSS--P-TFRPREILRCLQIGLLCVQERVEDRPMMSSVVLMLGSETALIPQPKQPGY     824
                * ** *    :*:* *: :*:******:** .*.** ***** ** **:  ***** * *

AlSRKb          CVGRSPFETDSSTHEQ--RDESCTVNQITISAIDPR     853
BrSRK9          WVRRSSYELDPSSSK--CDDDSWTVNQYTCSVIDAR     841
BrSRK46         CLIASYYANNPSSSRQFDDDESWTVNKYTCSVIDAR     860
BrSRK29         CVSGSSLETYSR-----RDDENWTVNQITMSIIDAR     854
BrSRK44         CVSGSSLETYSR-----RDDENWTVNQITMSIIDAR     854
BrSRK60         CVSQSSLETYSSWSK-LRDDENWTVNQITMSIIDAR     859
                 :  *              *:. ***: * * ** *
```

Supplemental Figure 4 Yamamoto et al.

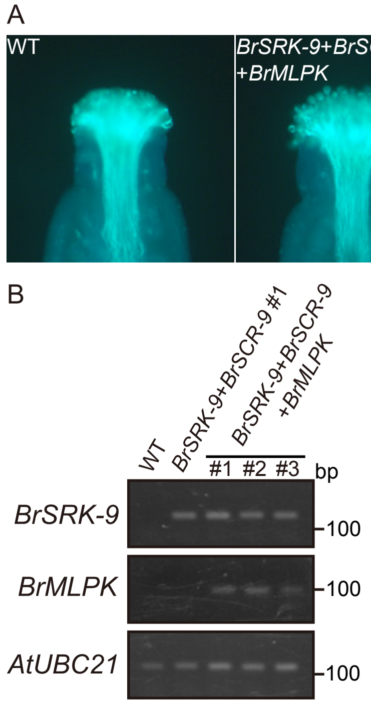


**Supplemental Figure 4.** *BrMLPK* does not affect the self-incompatibility (SI) response of *BrSRK-9+BrSCR-9* transgenic *A. thaliana* plants.

(A) Self-pollination assay of wild-type (WT) and transgenic *A. thaliana* plants expressing *BrSRK-9*, *BrSCR-9*, and *BrMLPK* (*BrSRK-9+BrSCR-9+BrMLPK*). A large number of self-pollen tubes were observed in the stigma of transgenic plants, similar to that observed in the stigma of wild-type plants. Scale bar = 100 μm.

(B) RT-PCR analysis of *BrSRK-9* and *BrMLPK* in the buds (positions -1 and -2) of wild-type (WT) plants, *BrSRK-9+BrSCR-9* transgenic *A. thaliana* line #1 (*BrSRK-9+BrSCR-9* #1), and *BrSRK-9+BrSCR-9+BrMLPK A. thaliana* lines #1, #2, and #3 (*BrSRK-9+BrSCR-9+BrMLPK* #1, #2, and #3). *AtUBC21* was used as an internal control.

Supplemental Figure 5 Yamamoto et al.

```
BrSRK9      RFLVSWRSSDDPSSGDFSYKLDIQRGLPEFYTFKDNTLVHRTGPWNGIRFSGIPEEQQLS    232
CgSRK7      RFLTSWKSSFDLSNGDYLFKLET-QGLPEFFLWKKFWILYRSGPWDGSRFSGMSEIQQWD    235
AlSRK25     RFLTCWKNSFDPSSGDYMFRLDT-QGLPEFFGLKNFLEVYRTGPWDGHRFSGIPEMQQWD    238
            ***..*:.* * *.**: ::*:  :*****:  *.hvI::*:***:* ****: * ** .

BrSRK9      YMVYNFTENSEEVAYTFLVTNNSIYSRLTINFSGFFERLTWTPSLVIWNPIWSSPASFQC    292
CgSRK7      DIIYNLTDNSEEVAFTFRLTDHNLYSRLTINDAGLLQQFTWDSTNQEWNMLWSTPK-EKC    294
AlSRK25     DIVYNFTENSEEVAYTFRLTDQTLYSRFTINSVGQLERFTWSPTQQEWNMFWSMPH-EEC    297
             ::**:*:******:** :*::.:***:***  * ::::**  :   ** :** *hvII:*

BrSRK9      DPYMICGPGSYCDVNTLPLCNCIQGFKPLNVQEWDMRDHTRGCIRRTRLSCRGDGFTRMK    352
CgSRK7      DYYDPCGPYAYCDMSTSPMCNCIEGFAPRNSQEWASGIVRGRCQRKTQLSCGGDRFIQLK    354
AlSRK25     DVYGTCGPYAYCDMSKSPACNCIKGFQPLNQQEWESGDESGRCRRKTRLNCRGDGFFKLM    357
            * *  *** :***:.. * ****:** * * ***  hvIII    * *:*:*.* ** * ::
```

**Supplemental Figure 5.** Amino acid sequence alignment of BrSRK9, *Capsella grandiflora* SRK7, and AlSRK25.

Amino acid sequences flanking the hv regions (outlined in red boxes) of BrSRK9, CgSRK7, and AlSRK25 were aligned using Clustal Omega. Amino acid residues essential and non-essential for SI are indicated in magenta and green, respectively.

**Supplemental Table 1.** qRT-PCR analysis of *Arabidopsis lyrata S-b* genes in transgenic *Arabidopsis thaliana* floral buds at different positions.

| Transgenic line ID | Gene name | Bud positions | Transcript levels[a] | | | Mean ± SD[b] |
|---|---|---|---|---|---|---|
| | | | Exp. 1 | Exp. 2 | Exp. 3 | |
| *AlSRK-b+AlSCR-b* #1 | *AlSRK-b* | +1, 0 | 0.65 | 0.83 | 0.63 | 0.70 ± 0.11[a] |
| | | -1, -2 | 1.00 | 1.00 | 1.00 | 1.00 ± 0.00[a] |
| | | -3, -4 | 0.79 | 1.38 | 0.99 | 1.05 ± 0.30[a] |
| | | -5, -6 | 0.58 | 0.92 | 0.62 | 0.71 ± 0.18[a] |
| | *AlSCR-b* | +1, 0 | 0.0062 | 0.015 | 0.0041 | 0.0085 ± 0.0059[a] |
| | | -1, -2 | 1.00 | 1.00 | 1.00 | 1.00 ± 0.00[ab] |
| | | -3, -4 | 2.35 | 6.92 | 2.27 | 3.84 ± 2.66[b] |
| | | -5, -6 | 0.76 | 1.44 | 0.66 | 0.95 ± 0.42[ab] |
| *AlSRK-b+AlSCR-b* #2 | *AlSRK-b* | +1, 0 | 1.09 | 0.46 | 1.05 | 0.86 ± 0.35[a] |
| | | -1, -2 | 1.00 | 1.00 | 1.00 | 1.00 ± 0.00[a] |
| | | -3, -4 | 0.88 | 1.80 | 0.80 | 1.16 ± 0.56[a] |
| | | -5, -6 | 1.03 | 0.93 | 0.85 | 0.94 ± 0.09[a] |
| | *AlSCR-b* | +1, 0 | 0.022 | 0.026 | 0.0058 | 0.018 ± 0.011[a] |
| | | -1, -2 | 1.00 | 1.00 | 1.00 | 1.00 ± 0.00[ab] |
| | | -3, -4 | 6.92 | 1.87 | 4.89 | 4.56 ± 2.54[b] |
| | | -5, -6 | 2.73 | 0.98 | 2.72 | 2.14 ± 1.01[ab] |

[a]Transcription levels of all genes were determined by the ΔΔCT method, with *AtUBC21* as the control. Relative expression levels in floral buds at positions -1 and -2 set at 1.

[b]Data represent mean ± standard deviation (SD) of three biological replicates (Exp. 1, 2, and 3). Different lowercase letters indicate statistically significant differences ($P < 0.05$; Tukey–Kramer method).

**Supplemental Table 2.** Absolute qRT-PCR analysis of transgenic *A. thaliana* plants expressing *AlSRK* and *Brassica rapa SRK* genes.

| Transgenic line ID | Relative transcript levels[a] | | | Mean ± SD[b] |
|---|---|---|---|---|
| | Exp. 1 | Exp. 2 | Exp. 3 | |
| *AlSRK-b+AlSCR-b* #1 | 1.82 | 0.84 | 0.48 | 1.04 ± 0.70[ac] |
| *AlSRK-b+AlSCR-b* #2 | 0.90 | 0.80 | 0.87 | 0.86 ± 0.05[ac] |
| *AlSRK-b+AlSCR-b* #13 | 0.80 | 0.62 | 0.93 | 0.78 ± 0.16[ac] |
| *BrSRK-9+BrSCR-9* #1 | 3.85 | 2.22 | 3.17 | 3.08 ± 0.82[d] |
| *BrSRK-9+BrSCR-9* #5 | 3.04 | 1.24 | 2.09 | 2.12 ± 0.90[cd] |
| *BrSRK-9+BrSCR-9* #6 | 6.98 | 4.81 | 6.15 | 5.98 ± 1.09[e] |
| *BrSRK-9 chimera+BrSCR-9* #2 | 0.08 | 0.17 | 0.09 | 0.11 ± 0.05[a] |
| *BrSRK-9 chimera+BrSCR-9* #3 | 1.41 | 1.23 | 1.40 | 1.35 ± 0.10[ac] |
| *BrSRK-9 chimera+BrSCR-9* #8 | 1.00 | 1.80 | 1.64 | 1.48 ± 0.42[ac] |
| *BrSRK-46 chimera+BrSCR-46* #2 | 0.98 | 0.93 | 0.64 | 0.85 ± 0.18[ac] |
| *BrSRK-46 chimera+BrSCR-46* #7 | 0.93 | 0.98 | 1.40 | 1.10 ± 0.26[ac] |
| *BrSRK-46 chimera+BrSCR-46* #10 | 1.34 | 1.88 | 0.88 | 1.37 ± 0.50[ac] |
| *BrSRK-29 chimera+BrSCR-29* #5 | 1.58 | 1.75 | 1.32 | 1.55 ± 0.22[bc] |
| *BrSRK-29 chimera+BrSCR-29* #11 | 0.10 | 0.14 | 0.13 | 0.12 ± 0.02[a] |
| *BrSRK-29 chimera+BrSCR-29* #16 | 0.17 | 0.09 | 0.09 | 0.12 ± 0.05[a] |
| *BrSRK-44 chimera+BrSCR-44* #2 | 0.28 | 1.05 | 0.65 | 0.66 ± 0.38[ab] |
| *BrSRK-44* chimera+*BrSCR-44* #3 | 0.60 | 0.19 | 0.23 | 0.34 ± 0.23[ab] |
| *BrSRK-44* chimera+*BrSCR-44* #4 | 0.42 | 0.98 | 1.05 | 0.82 ± 0.34[ac] |
| *BrSRK-60* chimera+*BrSCR-60* #3 | 0.76 | 0.52 | 0.48 | 0.59 ± 0.15[ab] |
| *BrSRK-60* chimera+*BrSCR-60* #5 | 0.40 | 0.24 | 0.36 | 0.33 ± 0.08[ab] |
| *BrSRK-60* chimera+*BrSCR-60* #7 | 0.89 | 0.62 | 0.72 | 0.74 ± 0.14[ac] |

[a]Relative transcript levels of all genes in floral buds at positions -1 and -2 were determined by the standard curve method, and *AtUBC21* transcript levels were set at 1.

[b]Data represent mean ± standard deviation (SD) of three biological replicates (Exp. 1, 2, and 3). Different lowercase letters indicate statistically significant differences ($P < 0.05$; Tukey–Kramer method).

**Supplemental Table 3.** Absolute qRT-PCR analysis of transgenic *A. thaliana* plants expressing *AlSCR* and *BrSCR* genes.

| Transgenic line ID | Relative transcript levels[a] | | | Mean ± SD[b] |
|---|---|---|---|---|
| | Exp. 1 | Exp. 2 | Exp. 3 | |
| *AlSRK-b+AlSCR-b* #1 | 7.04 | 13.41 | 9.78 | 10.08 ± 3.1[d] |
| *AlSRK-b+AlSCR-b* #2 | 10.21 | 7.57 | 7.36 | 8.38 ± 1.59[cd] |
| *AlSRK-b+AlSCR-b* #13 | 6.43 | 4.90 | 6.58 | 5.97 ± 0.93[c] |
| *BrSRK-9+BrSCR-9* #1 | 0.37 | 0.62 | 0.74 | 0.58 ± 0.19[ab] |
| *BrSRK-9+BrSCR-9* #5 | 0.18 | 0.19 | 0.17 | 0.18 ± 0.01[a] |
| *BrSRK-9+BrSCR-9* #6 | 1.69 | 1.86 | 1.59 | 1.72 ± 0.13[ab] |
| *BrSRK-9 chimera+BrSCR-9* #2 | 1.72 | 1.00 | 0.84 | 1.19 ± 0.47[ab] |
| *BrSRK-9 chimera+BrSCR-9* #3 | 0.29 | 0.60 | 0.30 | 0.40 ± 0.18[ab] |
| *BrSRK-9 chimera+BrSCR-9* #8 | 0.61 | 0.85 | 0.58 | 0.68 ± 0.15[ab] |
| *BrSRK-46 chimera+BrSCR-46* #2 | 1.35 | 0.70 | 0.71 | 0.92 ± 0.37[ab] |
| *BrSRK-46 chimera+BrSCR-46* #7 | 0.76 | 0.96 | 0.93 | 0.88 ± 0.11[ab] |
| *BrSRK-46 chimera+BrSCR-46* #10 | 0.57 | 0.70 | 0.78 | 0.68 ± 0.10[ab] |
| *BrSRK-29 chimera+BrSCR-29* #5 | 1.22 | 0.55 | 0.92 | 0.90 ± 0.34[ab] |
| *BrSRK-29 chimera+BrSCR-29* #11 | 0.25 | 0.22 | 0.55 | 0.34 ± 0.18[ab] |
| *BrSRK-29 chimera+BrSCR-29* #16 | 0.46 | 0.33 | 0.31 | 0.36 ± 0.08[ab] |
| *BrSRK-44 chimera+BrSCR-44* #2 | 1.01 | 1.13 | 1.30 | 1.15 ± 0.14[ab] |
| *BrSRK-44 chimera+BrSCR-44* #3 | 1.10 | 1.23 | 1.64 | 1.32 ± 0.28[ab] |
| *BrSRK-44 chimera+BrSCR-44* #4 | 1.10 | 0.55 | 0.64 | 0.76 ± 0.29[ab] |
| *BrSRK-60 chimera+BrSCR-60* #3 | 0.47 | 0.45 | 0.45 | 0.46 ± 0.01[ab] |
| *BrSRK-60 chimera+BrSCR-60* #5 | 3.16 | 2.62 | 2.69 | 2.82 ± 0.29[b] |
| *BrSRK-60 chimera+BrSCR-60* #7 | 0.46 | 0.97 | 0.31 | 0.58 ± 0.35[ab] |

[a]Relative transcript levels of all genes in floral buds at positions -3 and -4 were determined by the standard curve method, and *AtUBC21* transcript levels were set at 1.

[b]Data represent mean ± standard deviation (SD) of three biological replicates (Exp. 1, 2, and 3). Different lowercase letters indicate statistically significant differences ($P < 0.05$; Tukey–Kramer method).

**Supplemental Table 4.** List of primers used for plasmid construction.

| Primer name | Primer sequence (5'→3') |
|---|---|
| BrSRK9-F | AGAAAGTCGTGAAGGGTACCATGAAAGGTGTACGAAACAT |
| BrSRK9-R | TCCAAGGATCCCCGGTTAGCGGGCATCTATGACTG |
| BrSRK9(S)-R | TAGTAGCTTCTGTTTCCTTTTCCAGAGGCA |
| BrSRK29-F | AGAAAGTCGTGAAGGGTACCATGAAAAGGGTACAGAACAT |
| BrSRK29_44(S)-R | TAGTAGCTTCTGTTTCCTCCTCCAAAAGCA |
| BrSRK44-F | AGAAAGTCGTGAAGGGTACCATGAAAAGGGTAAAGAAC |
| BrSRK46-F | AGAAAGTCGTGAAGGGTACCATGAAAGGTGTACGAAACAT |
| BrSRK46(S)-R | TAGTAGCTTCTGTTTCCTTTTCCAAAGGCA |
| BrSRK60-F | AGAAAGTCGTGAAGGGTACCATGAAAGGGGTACATAACAT |
| BrSRK60(S)-R | TAGTAGCTTCTGTTTTCTCCTCCAAAAGCA |
| AlSRKb(kin)-F9 | TGCCTCTGGAAAAGGAAACAGAAGCTACTA |
| AlSRKb(kin)-F29_44 | TGCTTTTGGAGGAGGAAACAGAAGCTACTA |
| AlSRKb(kin)-F46 | TGCCTTTGGAAAAGGAAACAGAAGCTACTA |
| AlSRKb(kin)-F60 | TGCTTTTGGAGGAGAAAACAGAAGCTACTA |
| AlSRKb R | TCCAAGGATCCCCGGTTACCGAGGGTCGATGGCCG |
| BrSCR9-F | GCGGGTACCATGAAATCTGCTATTTATGCT |
| BrSCR9-R | CGCGTCGACTTATCTAACTTTGCATTTAAT |
| BrSCR46-F | GCGGGTACCATGAATTCTGCTATTTATGC |
| BrSCR46-R | GCGCTGCAGTTATTTACAATCGCAAGAAT |
| BrSCRII-F | GCGGGTACCATGAGATATGCTACTTCTATA |
| BrSCRII-R | GCGCTGCAGTTATGATTTAACTTTGCAAC |
| AlSCRbpro(H)-F | GGCCAGTGCCAAGCTATAGAAGAAAACCAGGCCTACATAGAACATGTTATA |
| AlSCRbpro(H)-R | GCAGGCATGCAAGCTGACAGATTTGATTGGTTAGT |
| AlSCRbpro(P)-F | CCAAGCTTGCATGCCATAGAAGAAAACCAGGCCTACATAGAACATGTTATA |
| AlSCRbpro(P)-R | GTTCTAGAGTCGACCGACAGATTTGATTGGTTAGT |
| AtS1-F | GCGCTGCAGATTCGAAATACATCGAGAA |
| AtS1-R | CGCTCTAGACTTCACGACTTTCTTTCTTAT |
| pRI201-StuI_F | GTTCTTCACTGTTGATAAGGCCTCCCGTCGATTAGCAAGT |
| pRI201-StuI_R | ACTTGCTAATCGACGGGAGGCCTTATCAACAGTGAAGAAC |

| | |
|---|---|
| M13_R | CAGGAAACAGCTATGAC |
| MLPK_InFusionF | TCACTGTTGATAAGGATGGGGATTTGCATGAGTGT |
| MLPK_InFusionR | CTAATCGACGGGAGGTCAGACAAACAGAGGCGAAG |
| BrSRK9(K206L)-F | TTCTATACATTTCTAGACAACACTCTAGTG |
| BrSRK9(K206L)-R | CACTAGAGTGTTGTCTAGAAATGTATAGAA |
| BrSRK9(V211E)-F | GACAACACTCTAGAGCATCGGACTGGTCCA |
| BrSRK9(V211E)-R | TGGACCAGTCCGATGCTCTAGAGTGTTGTC |
| BrSRK9(P282V)-F | GTGATATGGAACGTAATCTGGTCTTCTCCA |
| BrSRK9(P282V)-R | TGGAGAAGACCAGATTACGTTCCATATCAC |
| BrSRK9(P287I)-F | ATCTGGTCTTCTATAGCGAGCTTCCAGTGC |
| BrSRK9(P287I)-R | GCACTGGAAGCTCGCTATAGAAGACCAGAT |
| BrSRK9(F290H)-F | TCTCCAGCGAGCCACCAGTGCGATCCGTAC |
| BrSRK9(F290H)-R | GTACGGATCGCACTGGTGGCTCGCTGGAGA |
| BrSRK9(P294M)-F | TTCCAGTGCGATATGTACATGATTTGTGGG |
| BrSRK9(P294M)-R | CCCACAAATCATGTACATATCGCACTGGAA |
| BrSCR9(K55N)-F | TTCTATACGAATAATACGAATCAAAAGGCT |
| BrSCR9(K55N)-R | AGCCTTTTGATTCGTATTATTCGTATAGAA |
| BrSCR9(F69E)-F | TGTACAAGTCCTGAGCGAACTCGATATTGT |
| BrSCR9(F69E)-R | ACAATATCGAGTTCGCTCAGGACTTGTACA |
| BrSCR9(Y73L)-F | TTTCGAACTCGACTTTGTGATTGTGCAATT |
| BrSCR9(Y73L)-R | AATTGCACAATCACAAAGTCGAGTTCGAAA |

**Supplemental Table 5.** List of primers used for absolute qRT-PCR and reverse-transcription PCR (RT-PCR).

| Primer name | Primer sequence (5'→3') | Target gene |
|---|---|---|
| AlSRKb_kin RT_F | AATAACCTGCTCGGCTACGC | *AlSRK-b*, *BrSRK chimeras*, |
| AlSRKb_kin RT_R | CGAAACGCTGAAGGAGATGA | and *BrSRK-9 chimera* mutants |
| AlSCRb RT_F | GAACAAGTGCATGCGTTCTG | *AlSCR-b* |
| AlSCRb RT_R | TTGCACTGAATAGGCGTTCG | |
| BrSRK9 RT_F | CAATGTCGTCTGTGGTTTGGA | *BrSRK-9* |
| BrSRK9 RT_R | CAGGAATCATCGTCGCACTT | |
| BrSCR9 RT_F | CAAGAAGTGGAAGCTAATCTGAGG | Mutant *BrSCR-9* and *BrSCR-9* |
| BrSCR9 RT_R | AGAAGGCTTCGCAGTCATGC | |
| BrSCR29 RT_F | GGAGCTAGGAACTGCCCTGA | *BrSCR-29* |
| BrSCR29 RT_R | TCGGTCCGAAGTGTTTTTGAC | |
| BrSCR44 RT_F | CCGAATCGGGTCCTATCAGA | *BrSCR-44* |
| BrSCR44 RT_R | CCTCCCACGATTATGGTTGC | |
| BrSCR46 RT_F | AATCCGATGAAGCAGTGCAA | *BrSCR-46* |
| BrSCR46 RT_R | CTTACCCCGCGAACAAAAAT | |
| BrSCR60 RT_F | GGTGCATTAATTCCAGGAGCAC | *BrSCR-60* |
| BrSCR60 RT_R | ATCCTCGCAGGATTTTGCAT | |
| MLPK RT_F | CGACGTTGACAAAAACCAGAG | *BrMLPK* |
| MLPK RT_R | CGGTCTTATTTCCTCCTTCTTC | |
| UBC21 RT_F | AGAATGCTTGGAGTCCTGC | *AtUBC21* |
| UBC21 RT_R | AACCCTCTCACATCACCAGA | |
